# NGS data reveals the phyllosphere microbiome of wheat plants infected by the fungal pathogen *Zymoseptoria tritici*

**DOI:** 10.1101/2022.02.07.479402

**Authors:** Didac Barroso-Bergada, Marie Massot, Noémie Vignolles, Julie Faivre d’Arcier, Emilie Chancerel, Erwan Guichoux, Anne-Sophie Walker, Corinne Vacher, David A. Bohan, Valérie Laval, Frédéric Suffert

## Abstract

The fungal pathogen *Zymoseptoria tritici* is the causal agent of Septoria tritici blotch (STB), a major wheat disease in Western Europe. Microorganisms inhabiting wheat leaves might act as beneficial, biocontrol or facilitating agents that could limit or stimulate the development of *Z. tritici*. Improving our understanding of microbial communities in the wheat phyllosphere would lead to new insights into STB management. This resource announcement provides fungal and bacterial metabarcoding datasets obtained by sampling wheat leaves differing in the presence of symptoms caused by *Z. tritici*. Tissues were sampled from three wheat commercial varieties on three sampling dates during a cropping season. Weeds around wheat fields were sampled as well. In total, more than 450 leaf samples were collected. The pathogen *Z. tritici* was quantified using qPCR. We provide the raw metabarcoding datasets, the Amplicon Sequence Variant (ASV) tables obtained after bioinformatic processing, the metadata associated to each sample (sampling date, wheat variety and tissue health condition), a preliminary descriptive analysis of the data, and the code used for bioinformatic and descriptive statistical analysis.

## 1. Introduction

Wheat crops are exposed to several fungal plant pathogens, including *Zymoseptoria tritici*, the causal agent of Septoria tritici blotch (STB), a major disease in Western Europe (Fones and Gurr, 2015). In field conditions, wheat leaves host a multitude of other microorganisms — endophytic, epiphytic, pathogenic and saprophytic (Błaszczyk et al., 2021) — some of which interact directly or indirectly with *Z. tritici* (Kerdraon et al., 2019). Several taxa may also have antagonistic or synergistic activity while interacting with other taxa, and could be considered as potential biocontrol agents or facilitating agents that can limit or stimulate STB development (Chaudhry et al., 2020). Maximizing the chance of highlighting important interactions, for instance within cooccurrence network analysis, requires a thorough description of communities, under different biotic and abiotic conditions (Röttjers and Faust, 2018). In this resource announcement, we present fungal and bacterial community datasets collected on wheat leaves over the course of a wheat cropping season, taking into account: (i) the physiologic stage of wheat; (ii) the dynamics of STB development; and (iii) different wheat cultivars. We collected leaf samples in monovarietal plots grown with three wheat cultivars, at three dates over a growing season. One of the wheat varieties carried resistance genes to STB that also potentially impact the development of other constituent taxa of the microbial community. Three types of leaf sections were collected, which differed in the presence of symptoms caused by *Z. tritici*: (i) sections with no STB lesion from a visually healthy leaf; (ii) sections with no STB lesion from a visually symptomatic leaf; and (iii) sections with STB lesions. This dataset could be used to explore the cooccurrence of microbial species and thereby improve our understanding of the community dynamics associated with the development of *Z. tritici* on wheat leaves. Weeds in the margins and edges of cultivated wheat fields can act as alternative hosts for microbial species present in this crop. For this reason, a second dataset composed of weed leaf samples was collected to get an insight into the host range of the microorganisms associated to wheat leaves and potentially interacting with *Z. tritici* in the agrosystem.

## 2. Methods

### 2.1. Sampling

#### 2.1.1. Wheat

Samples were collected in 2018 at the Grignon experimental station (Yvelines, France; 48° 51’ N, 1° 58’ E) from three varieties of winter-sown, bread wheat (*Triticum aestivum*). Two varieties, Soissons (SOI) and Apache (APA), were considered susceptible to *Z. tritici* (both rated 5 on the ARVALIS-Institut du Végétal/CTPS scale, from 1 to 9, with 9 corresponding to the most resistant cultivar), while the variety Cellule (CEL), carrying the gene *Stb16q*, was considered to be more resistant (rated 7). Leaf samples of each variety were collected in three plots of 30 m^2^. The three APA and CEL plots were independent experimental plots described in Orellana-Torrejon et al. (2022) while the three SOI plots were delineated within a larger (1 ha) wheat field described in Morais et al. (2016) and Kerdraon et al. (2019). Within each plot, five samples were taken at locations spaced of 1 m apart along a transect. For each sample, three pieces of leaf were collected: an asymptomatic leaf piece taken from a leaf without any STB lesion (G); an asymptomatic leaf piece from a leaf with STB lesion (GS); and a symptomatic leaf piece including a portion of sporulating lesion (S), *i.e*. bearing pycnidia (*Z. tritici* asexual fruiting bodies). Three sampling campaigns were performed: the first on March 14th (SOI) and 15th (APA and CEL); the second on May 3rd; and the third on June 13th. In March and May, leaf pieces measured 5 cm long. G samples were taken from the central part of the second leaf (F2) of a plant located as close as possible to the sampling point. S and GS samples were collected from the third leaf (F3) of another plant. S samples were taken from the distal part of the leaf and GS samples were cut from the basal part (closer to the stem insertion) of the same leaf. In June, leaf pieces measured 3 cm long because leaves were broader and our goal was to collect an approximately similar amount of tissue on all sampling dates. All leaves were found to be symptomatic in June so we only collected S and GS samples from the third leaf of different plants.

#### 2.1.2. Weeds

On July 16th, samples were collected on eight species of weeds to produce a complementary dataset. Some of these weeds presented sparse symptoms, caused by undetermined fungal pathogens that had a very high probability of not being *Z. tritici*, specific to wheat. Five GS and five S samples were collected on *Lolium perenne* (LOLPE) individuals growing within the SOI field and on *Arrhenatherum elatius* (ARREL) individuals growing on a slope 5 m away from the SOI field. These two weed species were dominant weeds at the time of sampling. Five G samples were also collected on *Senecio vulgaris* (SENVU) individuals growing within the SOI field, *Poa annua* (POAAN) individuals growing on a path along the SOI field, *Hordeum murinum* (HORMU) and *Plantago lanceolata* (PLALA) individuals growing between the field and the path, and *Urtica dioica* (URTDI) and *Geranium molle* (GERMO) individuals growing on the slope 2 m away from the SOI field. All leaf samples were cut with scissors and placed in 2 mL autoclaved collection tubes. They were then brought back to the laboratory and stored at −20°C prior to freeze-drying.

### 2.2. DNA extraction

Total DNA was extracted with the DNeasy Plant Mini kit (Qiagen, France), using a protocol slightly modified from that recommended by Kerdraon et al. (2019). Two autoclaved DNAase-free inox 420C beads were added to each tube and samples were ground at 1500 rpm with the Geno/Grinder^®^ for 30 s, then twice 1 min, with manual shaking between each grinding step. Tubes were then centrifuged for 1 min at 6000 g. Leaf powder and 200 μL of buffer AP1 preheated to 60°C were mixed by vortexing the tubes for 30 s twice at 1500 g,and centrifuging them for 1 min at 3000 g. 250 μL of preheated buffer AP1 and 4.5 μL of RNase A were added to each tube and mixed by vortexing the tubes for 30s twice at 1500 g. After 5 min of rest, 130 μL of buffer P3 was added to each tube, which was then mixed by gentle inversion for 15 s, incubated at −20°C for 10 min and centrifuged for 1 min at 5000 g. The supernatant (450 μL) was transferred to a spin column and centrifuged for 2 min at 20000 g. The filtrate (200 μL) was transferred to a new tube, to which sodium acetate (200 μL, 3 M, pH 5) and cold 2-propanol (600 μL) were added. DNA was precipitated by incubation at −20°C for a minimum of 1 h and recovered by centrifugation (20 min, 13000 g). The pellet was washed with cold ethanol (70%), dried at 50°C for about 30min, and dissolved in 100 μL of AE buffer.

### 2.3. Bacterial 16S amplification

The V5-V6 region of the bacterial 16S rDNA gene was amplified using primers 799F-1115R (Redford et al., 2010; Chelius and Triplett, 2001) to exclude chloroplastic DNA. To avoid a two-stage PCR protocol and reduce PCR biases, each primer contained the Illumina adaptor sequence, a tag and a heterogeneity spacer, as described in Laforest-Lapointe et al. (2017) (799F: 5^*1*^-CAAGCAGAAGACGGCATACGAGATGTGACTGGAGTTCAGACGTGTGCTCTTCC GATCTxxxxxxxxxxxxHS-AACMGGATTAGATACCCKG-3^*1*^; 1115R:5^*1*^-AATGATACGGCGACCACCGAGATCTACACTCTTTCCCTACACGACGCTCTTCCG ATCTxxxxxxxxxxxxHS-AGGGTTGCGCTCGTTG-3^*1*^, where HS represents a 0-7-base-pair heterogeneity spacer and “x” a 12 nucleotide tag). The PCR mixture (20 μL of final volume) consisted of 4 μL of buffer Phusion High Fidelity 5X (ThermoFisher) (1X final), 2 μL each of the forward and reverse primers (0.2 μM final), 2 μL of 2 mM dNTPs (200 μM final), 8.2 μL of water, 0.6 μL of SO, DM0, 2 μL of Phusion Hot Start II Polymerase (ThermoFisher) and 1 μL of DNA template. PCR cycling reactions were conducted on a Veriti 96-well Thermal Cycler (Applied Biosystems) using the following conditions: initial denaturation at 98°C for 30s followed by 30 cycles at 98°C for 15 s, 60°C for 30 s, 72°C for 30 s with final extension of 72°C for 10 min. Two bacterial strains (*Vibrio splendidus* et *Sulfitobacter pontiacus*) were used as positive controls as they were unlikely to be found in our samples. The negative PCR controls were represented by PCR mix without any DNA template. Each PCR plate contained are negative extraction control, three negative PCR controls, one single-strain positive control and one two-strain positive control.

### 2.4. Fungal ITS amplification

The ITS1 region of the fungal ITS rDNA gene (Schoch et al., 2012) was amplified using primers ITS1F-ITS2 (White et al., 1990; Gardes and Bruns, 1993). To avoid a two-stage PCR protocol, each primer contained the Illumina adaptor sequence and a tag (ITS1F: *5^1^-* CAAGCAGAAGACGGCATACGAGATGTGACTGGAGTTCAGACGTGTGCTCTT CCGATCTxxxxxxxxxxxxCTTGGTCATTTAGAGGAAGTAA-3^*1*^; ITS2: 5^*1*^-AATGATACGGCGACCACCGAGATCTACACTCTTTCCCTACACGACGCTCTTCCG ATCTxxxxxxxxxxxxGCTGCGTTCTTCATCGATGC-3^*1*^, where “x” is the 12 nucleotide tag). The PCR mixture (20 μL of final volume) consisted of 10 μL of 2X QIAGEN Multiplex PCR Master Mix (2X final), 2 μL each of the forward and reverse primers (0.1 μM final), 4 μL of water, 1 μL of 10 ng*/*μL BSA and 1 μL of DNA template. PCR cycling reactions were conducted on a Veriti 96-well Thermal Cycler (Applied Biosystems) using the following conditions: initial denaturation at 95°C for 15 min followed by 35 cycles at 94°C for 30 s, 57°C for 90 s, 72°C for 90 s with final extension of 72°C for 10 min. ITS1 amplification was confirmed by electrophoresis on a 2% agarose gel. Two marine fungal strains (*Candida oceani* and *Yamadazyma barbieri*) were used as positive controls as they were unlikely to be found in our samples. One positive control included 1 μL of 10 ng*/*μL DNA of *Candida oceani* only and the other included an equimolar mixture of both strains. The negative PCR controls were represented by PCR mix without any DNA template. Each PCR plate contained one negative extraction control, three negative PCR controls, one single-strain positive control and one two-strain positive control.

### 2.5. Sequencing

MiSeq sequencing PCR products purification (CleanPCR, MokaScience), library sequencing on an Illumina MiSeq platform (v2 chemistry, 2 × 250 bp) and sequence demultiplexing (with exact index search) were performed at the PGTB sequencing facility (Genome Transcriptome Platform of Bordeaux, Pierroton, France). Fungal ITS1 amplicons were sequenced on three runs and bacterial 16S amplicons were sequenced on four runs.

### 2.6. Bioinformatic treatment

The MiSeq sequences produced were processed using the DADA2 pipeline version 1.22.0 (Callahan et al., 2016) and implemented in R. Primers were identified and removed using cutadapt 3.2 (Martin, 2011) and the trimmed sequences were then parsed to the DADA2 algorithm. Chimeras were removed using the removeBimeraDenovo functionality of DADA2. ASVs taxonomic assignment was performed using an implementation of the Naive Bayesian Classifier (Wang et al., 2007) included in the DADA2 pipeline. The databases used for taxonomic assignment were the Silva v138.1 (Quast et al., 2012) and the UNITE all eukaryotes v8.3 (Abarenkov et al., 2021) for 16S and ITS sequences, respectively. Three tables were obtained at the end of this process: an ASV table with the sequence count in each sample; a table with the taxonomic assignment of each ASV sequence; and a metadata table describing the collection conditions of each sample. The three tables were joined in a phyloseq object using the phyloseq bioconductor package v1.38.0 (McMurdie and Holmes, 2013). To filter out possible contaminants, the combined method of the isContaminant function of the DECONTAM Bioconductor package v1.14.0 (Davis et al., 2018) was used, followed by the decontamination method described in Galan et al. (2016). Moreover, 16S ASVs identified as chloroplastic or mitochondrial with Metaxa2.2.3 (Bengtsson-Palme et al., 2015), or according to their taxonomic assignment in the Silva database, were removed. The remaining ASVs were clustered using the Lulu algorithm (Frøslev et al., 2017) with default parameters. ASVs that could not be assigned to a bacterial or fungal phylum were removed. Finally, ASVs present in less than 1% of the samples were removed to make sure that the data were free of sequencing artifacts and low abundant contaminants (Cao et al., 2021).

### 2.8. Quantification of *Z. tritici* by qPCR

The abundance of *Z. tritici* in wheat tissues was estimated using the quantitative PCR assay developed by Duvivier et al. (2013). The specific set of primers included a forward primer (5’-ATTGGCGAGAGGGATGAAGG-3’), a reverse primer (5’-TTCGTGTCCCAGTGCGTGTA-3’), both leading to an amplification product of 101 pb, and a Taqman fluorogenic probe (5’-ACGACTCGCGGCTTTCACCCAACG-3’). The probe was labelled with a FAM fluorescent reporter dye and a BHQ-1 quencher. The quantification reaction was performed with the CFX96 Real time System C1000 Thermal Cycler (BIORAD, USA), using hard shell PCR 96-well WHT/CLR plates. The mix reaction was composed of reverse and forward primers at 500 nM per reaction, the probe at 500 nM per reaction in a final volume of 25 μL, with 5 μL of DNA introduced per well. All samples (standard DNA, eDNA to be analyzed, and negative controls) were analyzed with three replicates. The PCR program was 95°C for 10 min and (95°C for 15 s, 60°C for 20 s, 72°C for 40 s) repeated for 40 cycles. The concentration of DNA in the unknown samples was calculated by comparing cycle threshold (Ct) values of the samples with known standard quantities of *Z. tritici* genomic DNA, using a tenfold serial dilution from 5 ng to 5.10^-3^ ng per well. Ct values were plotted against the log of the initial concentration of *Z. tritici* genomic DNA to produce the standard curve used for sample quantity determination.

### 2.9. Analysis

Data contained in the phyloseq object were analyzed using the statistic environment R v4.1.2 (R Core Team, 2020) to characterize the fungal and bacterial community composition and to assess the effect of the different experimental factors on these communities. The analysis was performed using only the samples obtained from wheat plants at March and May. Samples obtained in June were not included in the analysis because there were no healthy (G) samples available. ASV counts were transformed using a clr transformation (Aitchison, 1982) to obtain scale-invariant values, avoiding the compositional effect introduced during the sequencing process (Gloor et al., 2017). Then, the phyloseq R package was used to obtain the euclidean distance between samples and to perform a Principal Coordinate Analysis (PCoA). The PCoA was plotted using ggplot2 package v3.3.5 (Wickham, 2016). A permutational multivariate analysis of variance (permanova), performed to assess the effect of the experimental design on the communities, was done using the adonis2 function of the vegan R package v2.5.7(Oksanen et al., 2019) following the experimental formula “tissue × date × variety/plot”. Alpha diversity measures were obtained using the phyloseq package and fitted in a generalised mixed model using the lme4 R package v1.1-27.1 (Bates et al., 2015). *Z. tritici* qPCR analysis was also fitted in a generalised mixed model using lme4.

## 3. Results

This resource announcement provides two sets of raw sequences files, one set obtained using primers for the fungal ITS region and another obtained using primers for the bacterial 16S region. The sequences are available in the Dataverse files (see section Availability of Data and Materials). The raw and filtered ASV tables obtained during the dereplication and filtering process are provided in the form of phyloseq objects (McMurdie and Holmes, 2013). Each phyloseq object includes the ASV table, a table with the ASV taxonomic assignation and a metadata table. The raw ASV tables also include the positives and negatives control samples used for the filtering. The samples obtained in June, as well as samples obtained from weeds growing in the vicinity of the wheat crop, are included in the phyloseq objects but were not analyzed in the present study. The metadata table includes, for each sample, the wheat variety or weed species sampled, the sampling date, the plot and the wheat cultivar, the visual assessment of symptoms and the *Z. tritici* DNA concentration obtained by qPCR (Figure 1A, Table 1, Suppl. Table S1). The tables showing the change in number of reads in each sample during the bioinformatic process are also provided in the Dataverse files.

**Figure 1.**
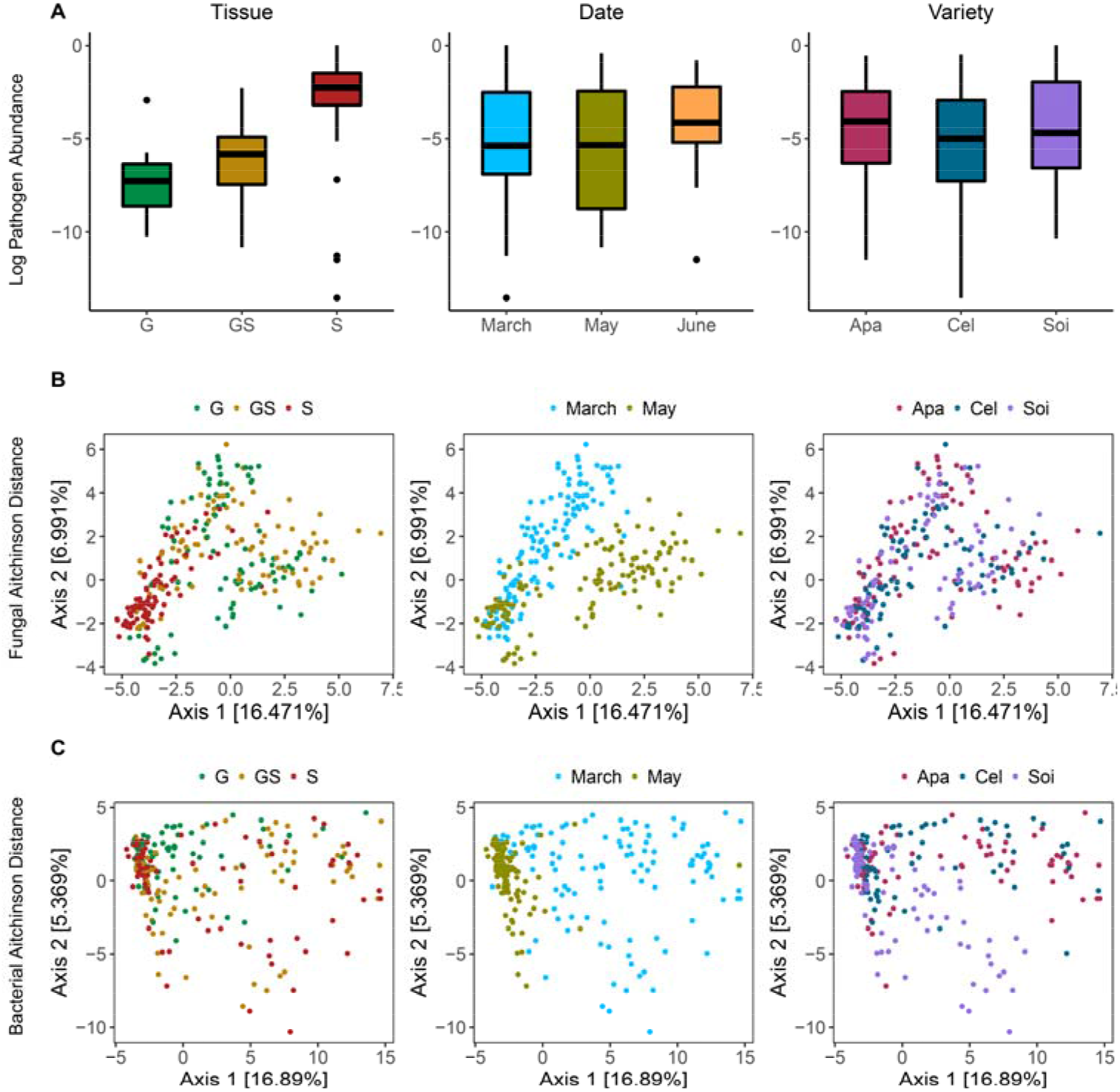
Community composition of wheat leaves. **A.** Abundance of the pathogen (*Zymoseptoria tritici*) on wheat leaves measured using qPCR. Community composition is represented as the relative abundance of each taxonomic unit for each experimental factor, at the genus level for fungi communities and order level for bacterial. **B, C.** Principal Coordinates Analysis plots showing the similarity of fungal (B) and bacterial (C) communities from different samples. The ordination was performed using the Aitchinson distance (Aitchison, 1982). Plots were colored by: the Septoria tritici blotch symptoms, G corresponding to leaf samples collected on asymptomatic leaves, GS corresponding to green parts of a symptomatic leaf, and S to symptomatic parts of a leaf; the sampling seasons, March and May; and the wheat varieties sampled, Apache (APA), Cellule (CEL), and Soissons (SOI).

**Table 1.**
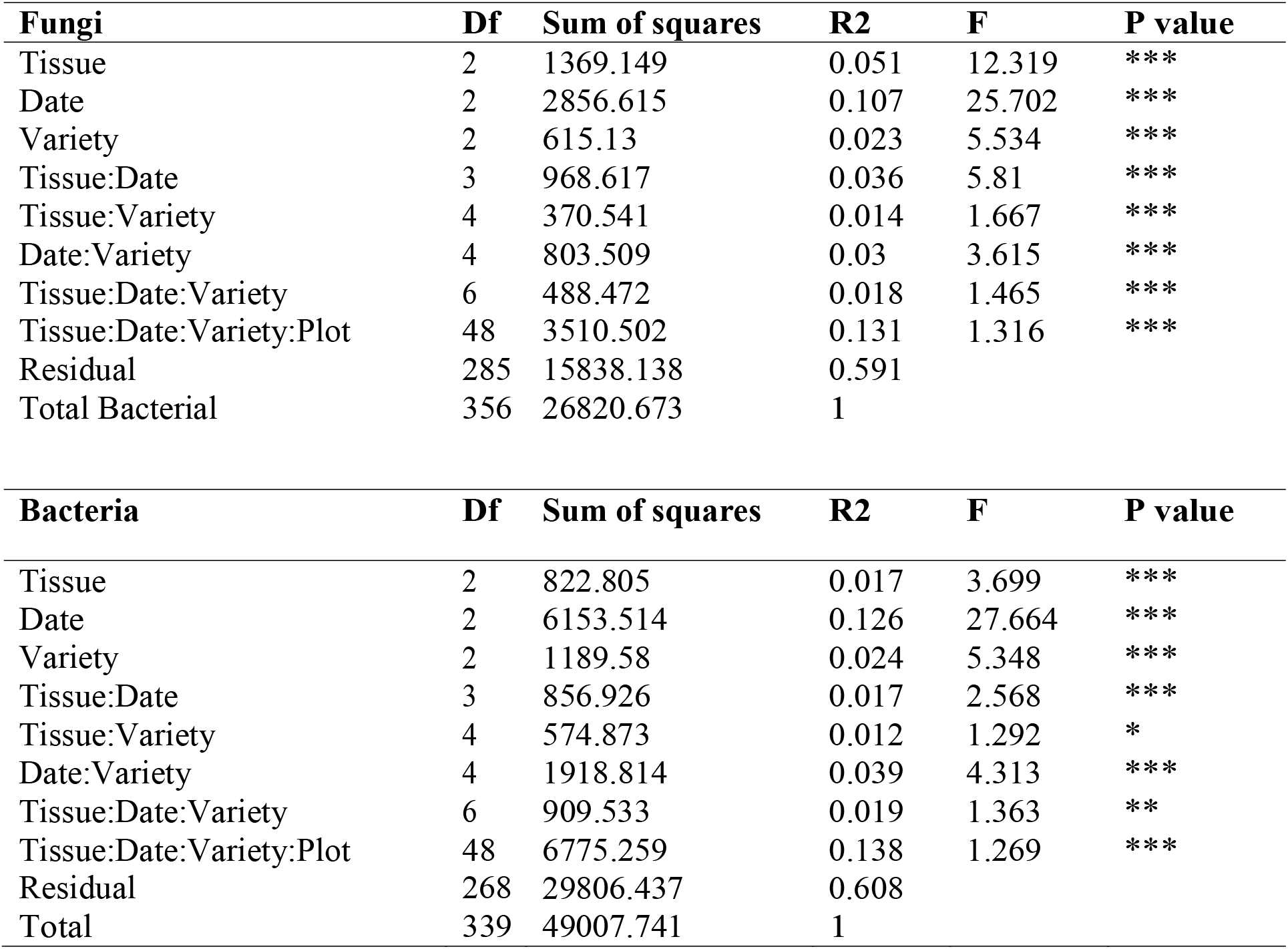
Permutational multivariate analysis of variance between samples. The Aitchison distance (Aitchison, 1982) between samples in the abundance matrix is used as a distance metric. The analysis was performed for the fungal and bacterial ASV tables separately using the R vegan package (Oksanen et al., 2019). Factors were: Tissue, corresponding to Septoria tritici blotch symptomatology; Date, corresponding to the sampling dates; and Variety, corresponding to the wheat varieties sampled. The plots were considered nested to the wheat variety. Values of p = * < .05; ** < .01; *** < .001.

### 3.1. Fungal communities

360 samples were sequenced using ITS primers and gave an average of 31,586 raw fungal sequences per sample with a minimum of 45 reads and a maximum of 246,434 reads per sample. The amplicon sequence variant (ASV) inference process identified an average of 28,178 high quality sequences per sample distributed in 2821 unique ASVs in the 360 samples. The ASV table obtained after the filtering process, which deleted contaminants and low abundant ASVs, was made up of an average of 27,609 sequences per sample distributed between 391 ASVs and 357 samples. 3 samples did not have any sequence after the filtering process. The minimum number of reads in a sample was 20 and the maximum was 223,756. 101 samples of weeds growing close to the field were also sequenced using ITS primers. The bioinformatic process allowed to obtain a mean of 45,631 sequences per sample and a total of 337 ASVs from these weed samples. The minimum number of reads in a sample was 1331 and the maximum was 365,035. The number of reads in each sample at each step of the bioinformatic process is supplied in the Dataverse files.

#### Taxonomic composition

For the wheat dataset, sequences assigned to the Ascomycota represented 74% of the total counts, while sequences assigned to Basidiomycota represented 25% (Figure 2A). As expected, sequences assigned to the genus *Zymoseptoria* were the most abundant (60% of the sequences). *Zymoseptoria* was also more abundant in symptomatic than in asymptomatic leaf samples. It was also slightly more abundant in the Soissons (SOI) and the Apache (APA) varieties, than in the Cellule (CEL) cultivar, which is less susceptible because it carries the *Stb16q* resistance gene.

**Figure 2.**
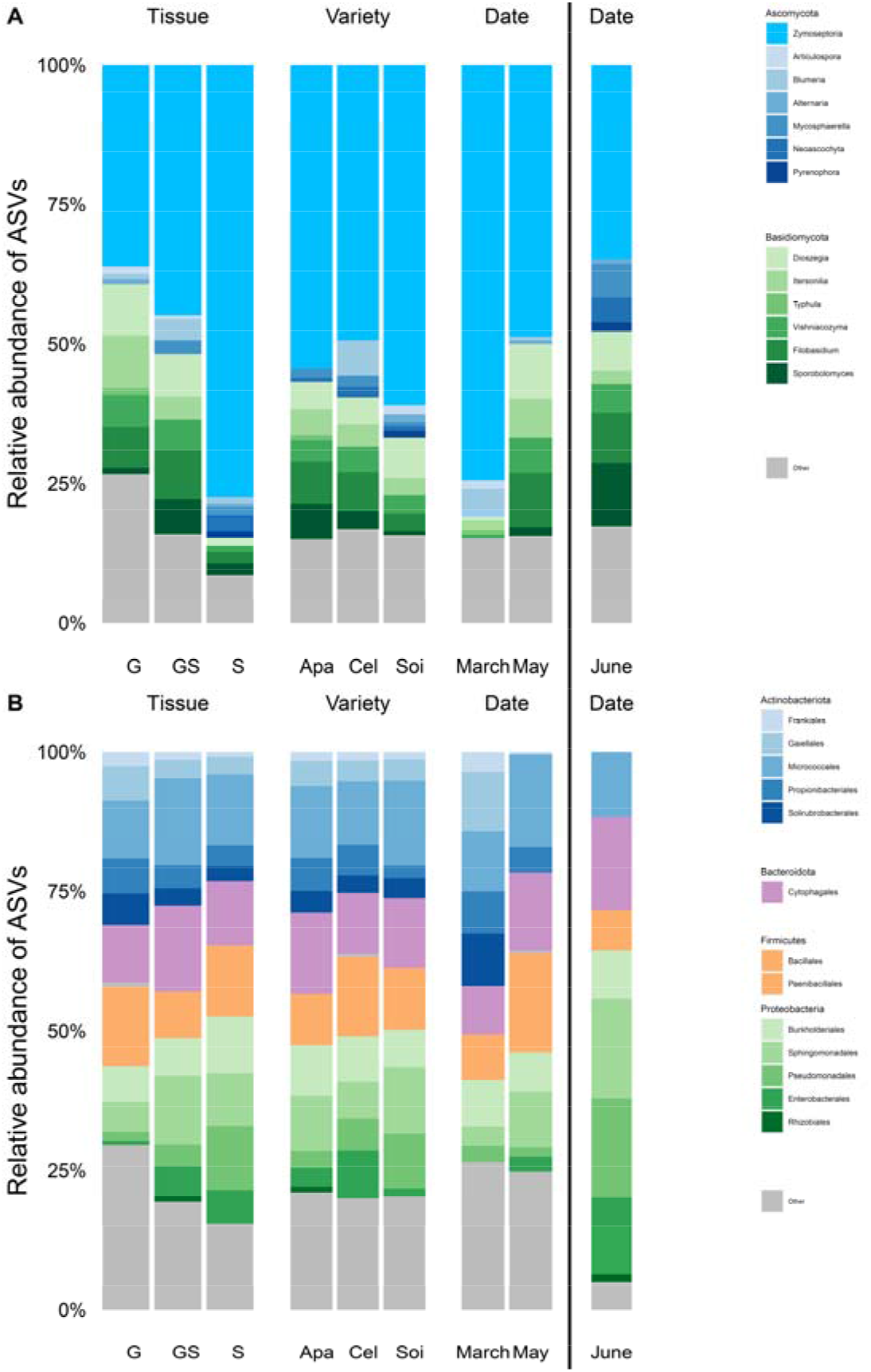
Barplots showing the relative abundance of different taxons along different dates, wheat varieties and leaves tissue conditions. Barplot **A** shows the relative abundance of different fungal genus. Barplot **B** shows the relative abundance of different bacterial orders. Barplots are separated by: Septoria leaf blotch symptoms, G corresponding to leaf samples collected on asymptomatic leaves, GS corresponding to green parts of a symptomatic leaf and S to symptomatic part of a leaf; the sampling seasons, March, May and June (June is separated by a bar because no asymptomatic samples (G) where available).

#### Alpha diversity

Fungal community richness, evenness and diversity differed significantly between dates and tissue health conditions (Suppl. Figure S1, Suppl. Table S2). Wheat variety had minor effect in all alpha diversity measures, being significant only for richness and diversity.

#### Beta diversity

The composition of wheat foliar fungal communities differed significantly among dates, varieties and tissue health conditions (Figure 1B, Suppl. Figure S1, Table 1, Suppl. Table S2). Tissue was the most important factor, explaining 10% of the variance. Samples collected in June were not included in the permutational analysis of variance to avoid apotential bias caused by the absence of healthy leaves (G samples) at that time.

### 3.2. Bacterial communities

The 360 samples used for ITS sequencing were also sequenced using 16S primers, obtaining an average of 40,724 raw bacterial sequences per sample with a minimum of 0 read and a maximum of 92520 reads per sample. The amplicon sequence variant (ASV) inference process identified a mean of 31,969 high quality sequences shared between 12,349 unique ASVs in 350 samples. 10 samples did not generate enough sequences to perform the ASV inference. The ASV table obtained after the filtering process, carried out to delete contaminants and low abundant ASVs, was composed of an average of 13,964 sequences per sample distributed between 1495 ASVs and 340 samples. The minimum number of reads in a sample was 2 and the maximum was 71,051. 102 samples of weeds growing surrounding the wheat plots were also sequenced using 16S primers. The bioinformatic process produced an average of 29,991 sequences per sample and 1,068 unique ASVs from the weed samples. The minimum number of reads in a sample was 30 and the maximum was 70,999.The number of reads of each sample at each step of the bioinformatic process is supplied in the Dataverse files.

#### Taxonomic composition

The most abundant bacterial phyla were Proteobacteria (39% of sequence counts), followed by Actinobacteria (35%), Bacteroidetes (12%) and Firmicutes (11%) (Figure 2B). Proteobacteria were more evident in later sampling dates while Actinobacteria were more present at the March sampling. Tissue condition and wheat variety did not seem to have an important effect on the community composition of the phyla.

#### Alpha diversity

Bacterial community richness, evenness and diversity differed significantly between tissue health conditions (Suppl. Figure S1, Suppl. Table S3). On the other side, sampling date had effect on richness and evenness measures, while wheat variety only had effect on the diversity measure.

#### Beta diversity

The composition of the bacterial communities of wheat leaves differed significantly among dates, varieties and tissue conditions (Figure 1C, Suppl. Figure S1, Table 1, Suppl. Table S3). As with the fungal component of the community, date was the most important structuring factor, explaining some 8% of the variance in the bacterial community (Table 1). Samples obtained in June were not included in the permutational analysis to avoid a potential for bias caused by the absence of healthy leave samples on that sample date.

## 4. Conclusions

Preliminary statistical analyses revealed that sampling date, wheat variety and STB symptoms had significant effects on fungal and bacterial communities of the wheat phyllosphere. While the three factors tested structured the community, the date of sampling exhibited the strongest effect. As expected, we found congruence between the presence of *Z. tritici*, assessed by eye (STB symptoms on the leaves) and by qPCR (concentration of *Z. tritici* DNA within the leaf tissues). This finding confirms the relevance of the sampling strategy to meet the assigned objectives. Co-occurrence network analyses could be used to characterize the dynamics of the community associated with *Z. tritici* and might help identify individual taxa of interest as potential biocontrol or beneficial agents to improve “wheat health”. The weed dataset could also be examined for *Z. tritici* interactions with different communities present on non-crop plants within and in the margins of the field.

## Availability of Data and Materials

The sequence datasets were deposited in NCBI SRA in bioproject PRJNA803042 (https://www.ncbi.nlm.nih.gov/bioproject/803042). The biosample accession numbers are SAMN25610777 to SAMN25611238. Bioinformatic scripts and raw and filtered ASV tables in R phyloseq format were deposited in Dataverse (https://doi.org/10.15454/QTXFP9). The tables showing variation in sequence counts during the bioinformatic process and the scripts used for data processing and statistical analysis were included in the Dataverse deposit.

## Author contributions

Author contributions were as follows: (i) D.A.B., C.V., F.S. and V.L coordinated the study; (ii) F.S., V.L., A.-S.W. and M.M. designed the sampling, collected samples and metadata; (iii) N.V., J.F.A., E.C. and E.G. developed protocols and performed the molecular biology work; (iv) D.B.B. performed data analysis and wrote the manuscript; (v) all authors edited the manuscript.

## Acknowledgments

We thank all members of the *Consortium Biocontrôle* for their support for the project. We thank Céline Lalanne, Adline Delcamp, Christophe Boury and all other members of PGTB (Genome Transcriptome Platform of Bordeaux) for their support in molecular biology, and Gaétan Burgaud and Frédéric Garabetian for kindly providing marine strains used as positive controls. We also thank Frédéric Barraquand, Stéphane Robin, Charlie Pauvert and Andreas Makiola for helpful discussions at the beginning of the project. We acknowledge support from the ANR NGB project (ANR-17-CE32-0011). PTGB and BIOGER received support from *Investissements d’avenir* and *convention attributive d’aide EquipEx Xyloforest* (ANR-10-EQPX-16-01) and from *Saclay Plant Sciences-SPS* (ANR-17-EUR-0007), respectively.

---

**Suppl. Table S1.** Analysis of Variance (ANOVA) of the fitted linear mixed model of the abundance of *Zymoseptoria tritici*. *Z. tritici* abundance was obtained by qPCR. Values of p = * < .05; ** < .01; *** < .001.

|  | Chisq | Df | Pr..Chisq. |
| --- | --- | --- | --- |
| Tissue | 201.608 | 2 | *** |
| Variety | 18.065 | 2 | *** |
| Date | 13.743 | 2 | ** |
| Tissue:Variety | 37.47 | 4 | *** |
| Tissue:Date | 20.069 | 3 | *** |
| Variety:Date | 30.747 | 4 | *** |
| Tissue:Variety:Date | 35.217 | 6 | *** |

**Suppl. Table S2.** General Mixed models of three alpha diversity fungal community measures. Observed richness, Shannon diversity index and inverse Simpson evenness index are modelled. Richness is modeled using a Negative Binomial distribution while Diversity and Evenness are modelled using a Gaussian distribution. Fixed factors are: Septoria leaf blotch symptoms, G corresponding to green parts of asymptomatic leaves, GS corresponding to green parts of symptomatic leaves and S to symptomatic part of symptomatic leaves; the sampling seasons, March and May; and the wheat varieties sampled, Apache (APA), Cellule (CEL), and Soissons (SOI). Plot is considered a random factor. Values of p = * < .05; ** < .01; *** < .001.

|  |  |  |  |  |
| --- | --- | --- | --- | --- |
| Richness |  |  |  |  |
| Fixed-effects |  |  |  |  |
|  | Estimate | Std-error | z | Pval |
| (Intercept) | -5.837 | 0.144 | -40.611 | *** |
| VarietyCel | 0.222 | 0.133 | 1.676 | . |
| VarietySoi | 0.306 | 0.136 | 2.247 | * |
| DateMay | 0.417 | 0.113 | 3.705 | *** |
| TissueGS | -0.825 | 0.127 | -6.476 | *** |
| TissueS | -2.338 | 0.14 | -16.742 | *** |
| Random-effects |  |  |  |  |
| Group | Name | Variance | Std-Dev |  |
| Plot | Intercept | 0.003 | 0.052 |  |
| Diversity |  |  |  |  |
| Fixed-effects |  |  |  |  |
|  | Estimate | Std-error | z | Pval |
| (Intercept) | -10.324 | 0.487 | -21.182 | *** |
| VarietyCel | 0.791 | 0.573 | 1.38 |  |
| VarietySoi | 1.25 | 0.573 | 2.18 | * |
| DateMay | 2.156 | 0.273 | 7.887 | *** |
| TissueGS | -1.971 | 0.206 | -9.557 | *** |
| TissueS | -3.217 | 0.595 | -5.406 | *** |
| Random-effects |  |  |  |  |
| Group | Name | Variance | Std-Dev |  |
| Plot | Intercept | 6.948 | 2.636 |  |
| Evenness |  |  |  |  |
| Fixed-effects |  |  |  |  |
|  | Estimate | Std-error | z | Pval |
| (Intercept) | -11.708 | 0.408 | -28.669 | *** |
| VarietyCel | 0.388 | 0.534 | 0.726 |  |
| VarietySoi | 0.78 | 0.545 | 1.43 |  |
| DateMay | 1.063 | 0.183 | 5.802 | *** |
| TissueGS | -1.258 | 0.202 | -6.217 | *** |
| TissueS | -2.8 | 0.629 | -4.453 | *** |
| Random-effects |  |  |  |  |
| Group | Name | Variance | Std-Dev |  |
| Plot | Intercept | 0.117 | 0.342 |  |

**Suppl. Table S3.** General Mixed models of three alpha diversity bacterial community measures. Observed richness, Shannon diversity index and inverse Simpson evenness index are modelled. Richness is modeled using a Negative Binomial distribution while Diversity and Evenness are modelled using a Gaussian distribution. Fixed factors are: Septoria leaf blotch symptoms, G corresponding to green parts of asymptomatic leaves, GS corresponding to green parts of symptomatic leaves and S to symptomatic part of symptomatic leaves; the sampling seasons, March and May; and the wheat varieties sampled, Apache (APA), Cellule (CEL), and Soissons (SOI). Plot is considered a random factor. Values of p = * < .05; ** < .01; *** < .001.

|  |  |  |  |  |
| --- | --- | --- | --- | --- |
| Richness |  |  |  |  |
| Fixed-effects |  |  |  |  |
|  | Estimate | Std-error | z | Pval |
| (Intercept) | -3.621 | 0.113 | -31.979 | *** |
| VarietyCel | 0.243 | 0.134 | 1.812 | . |
| VarietySoi | 0.098 | 0.13 | 0.75 |  |
| DateMay | 0.16 | 0.078 | 2.059 | * |
| TissueGS | -0.66 | 0.093 | -7.118 | *** |
| TissueS | -0.673 | 0.097 | -6.936 | *** |
| Random-effects |  |  |  |  |
| Group | Name | Variance | Std-Dev |  |
| Plot | Intercept | 0.013 | 0.113 |  |
| Diversity |  |  |  |  |
| Fixed-effects |  |  |  |  |
|  | Estimate | Std-error | z | Pval |
| (Intercept) | -5.376 | 0.077 | -69.799 | *** |
| VarietyCel | 0.014 | 0.092 | 0.152 |  |
| VarietySoi | -0.532 | 0.164 | -3.239 | ** |
| DateMay | -0.219 | 0.163 | -1.348 |  |
| TissueGS | -0.462 | 0.101 | -4.587 | *** |
| TissueS | -1.116 | 0.139 | -8.011 | *** |
| Random-effects |  |  |  |  |
| Group | Name | Variance | Std-Dev |  |
| Plot | Intercept | 0 | 0 |  |
| Evenness |  |  |  |  |
| Fixed-effects |  |  |  |  |
|  | Estimate | Std-error | z | Pval |
| (Intercept) | -9.634 | 0.203 | -47.413 | *** |
| VarietyCel | 0.073 | 0.269 | 0.272 |  |
| VarietySoi | 0.398 | 0.271 | 1.47 |  |
| DateMay | 0.946 | 0.134 | 7.035 | *** |
| TissueGS | -1.051 | 0.142 | -7.384 | *** |
| TissueS | -1.567 | 0.168 | -9.32 | *** |
| Random-effects |  |  |  |  |
| Group | Name | Variance | Std-Dev |  |
| Plot | Intercept | 0.036 | 0.191 |  |

**Supplementary figure S1.**
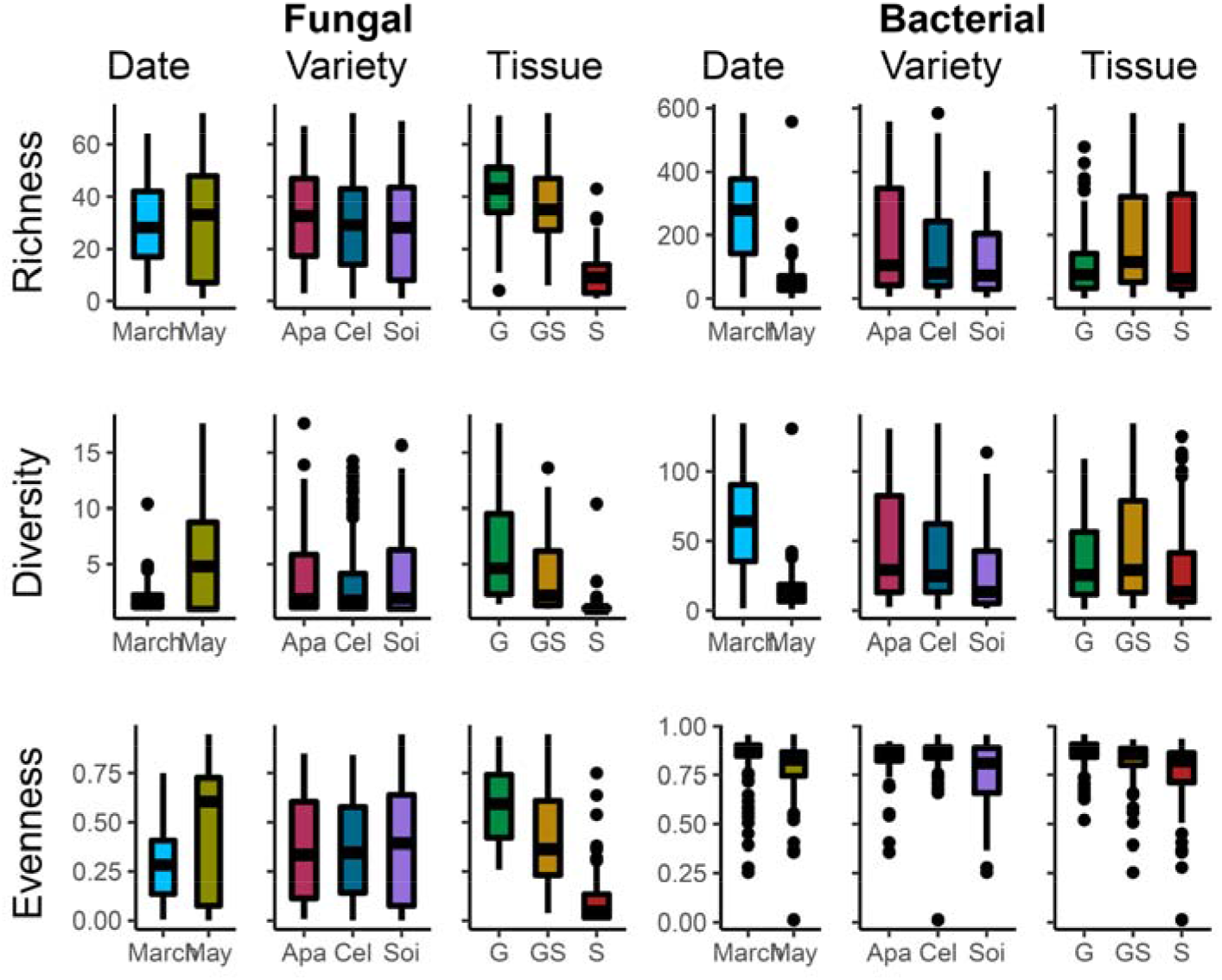
Boxplot showing three alpha diversity community measures for fungal and bacterial data. Observed richness, Shannon diversity index and inverse Simpson evenness index are plotted. Boxplots are separated by: Septoria tritici blotch symptoms, G corresponding to green parts of asymptomatic leaves, GS corresponding to green parts of symptomatic leaves and S to symptomatic part of symptomatic leaves; the sampling seasons, March and May; and the wheat varieties sampled, Apache (APA), Cellule (CEL), and Soissons (SOI). Plot was treated as a random factor.

